# Methanogens are associated with altered microbial production of short-chain fatty acids and human-host metabolizable energy

**DOI:** 10.1101/2024.12.31.630929

**Authors:** Blake Dirks, Taylor L. Davis, Elvis A. Carnero, Karen D. Corbin, Steven R. Smith, Bruce E. Rittmann, Rosa Krajmalnik-Brown

## Abstract

Methanogens are CH_4_-producing, H_2_-oxidizing (i.e. hydrogenotrophic) archaea. Numerous studies have associated methanogens with obesity, but these results have been inconsistent. One link to host metabolism may be methanogens’ ability to consume H_2_, thus reducing H_2_ partial pressure and thermodynamically enhancing fermentation of sugars to short-chain fatty acids (SCFA) that the host can absorb. Because research linking methanogenesis to human metabolism is limited, our goal with this exploratory analysis was to investigate relationships between methanogens and other hydrogenotrophs, and the association of methanogens with human metabolizable energy (ME). Using results from a randomized crossover feeding study with well-characterized human participants and novel continuous methane measurements, we analyzed hydrogenotroph abundance and activity, fecal and serum SCFAs, and host ME between high and low CH_4_ producers. Methanogens were detected in about one-half of participants, and most were high CH_4_ producers. We found no evidence that methanogens’ consumption of H_2_ to produce CH_4_ affected other hydrogenotrophs. High CH_4_ producers had greater serum propionate and host ME irrespective of diet, as well as greater gene and transcript abundance of a key enzyme of the H_2_-consuming, propionate-producing succinate pathway. Finally, a network analysis revealed positive relationships between *Methanobrevibacter smithii* (the most prevalent methanogen in the human colon) and bacteria capable of degrading fiber and fermenting fiber-degradation products, thus forming a trophic chain to extract additional energy from undigested substrates. Our results show that methanogens in a microbial consortium were linked to host metabolizable energy through enhanced SCFA production and host SCFA absorption.

## INTRODUCTION

The human body hosts around 38 trillion bacterial cells, most of which reside in the colon [1]. Besides bacteria, the intestinal microbiota also includes archaea, protists, and viruses [2]. One of the important functions of the intestinal microbiota is fermentation [3]. Unmetabolized macronutrients entering the colon become substrates for bacterial hydrolysis and fermentation, which generates simple products, such as H_2_, CO_2_, and short-chain fatty acids (SCFAs) [4].

While SCFAs have received much attention due to their impact on host health [5], H_2_ remains understudied despite its importance to the microbiome. High H_2_ partial pressure inhibits bacteria from regenerating NAD^+^ from NADH, which decreases the fermentation of complex substrates and impairs microbial growth [6,7]. H_2_ partial pressure also influences the thermodynamics of SCFA production: Low partial pressure favors acetate and butyrate production, while high partial pressure favors propionate production [8].

The H_2_ produced by fermentation can be oxidized by microbes known as hydrogenotrophs in well-known microbial processes that produce acetate, sulfide, or methane [9]. The three common groups of hydrogenotrophic microorganisms in the human colon are homoacetogenic bacteria, sulfate-reducing bacteria (SRB), and methanogenic archaea. These three groups can use H_2_ as an electron donor to produce energy [9]. **Figure 1** summarizes the hydrogenotrophic groups, their substrates and products, and the fate of their products.

**Figure 1.**
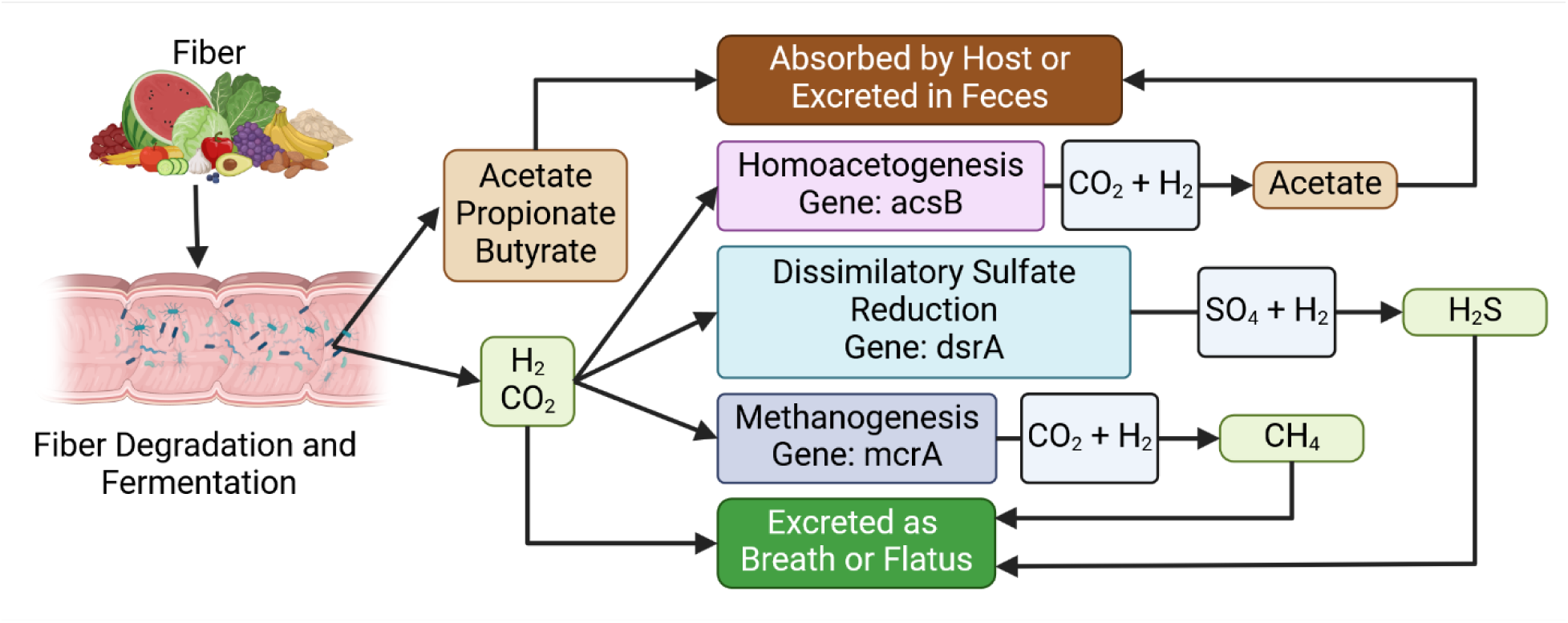
Hydrogenotrophic processes in the human colon when the macronutrient input is carbohydrate. Fiber that reaches the colon is hydrolyzed and fermented by the microbiome to produce SCFAs, H_2_, and CO_2_. Most SCFAs are absorbed by the host or excreted in feces. While some H_2_ is released through the breath or flatus, most is metabolized in the colon by homoacetogenesis, dissimilatory sulfate reduction, or methanogenesis. Acetate produced though hydrogenotrophy is absorbed by the host or excreted in feces. CH_4_ and some hydrogen sulfide produced through hydrogenotrophy are released through the breath or flatus. Figure created using Biorender.

Homoacetogenic bacteria, which include species in the genera *Blautia*, *Clostidium*, and *Ruminococcus*, oxidize H_2_ and reduce CO_2_ to make acetate using the Wood-Ljungdahl pathway [10]. The stoichiometry for homoacetogenesis is:

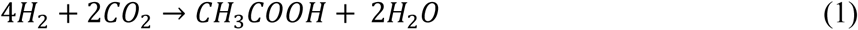

Homoacetogens carrying out reaction (1) are autotrophs, which means that their carbon source is inorganic carbon, or CO_2_. However, most homoacetogens are not obligate hydrogenotrophs and can ferment organic substrates, such as glucose, as the electron donor and carbon source [11].

SRB in the genera *Desulfovibrio* and *Fusobacteria* oxidize H_2_ and reduce sulfate to sulfide through dissimilatory sulfate reduction [12]:

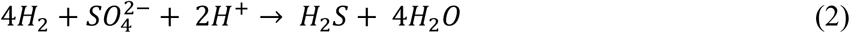

Like homoacetogens, SRB can also ferment organic substrates, such as lactate and pyruvate [11]. The predominant methanogen in the human colon is *Methanobrevibacter smithii*, although *Methanosphaera stadtmanae* and *Methanomassiliicoccus* spp are sometimes detected [13]. Methanogens can be categorized into three groups: 1) hydrogenotrophic methanogens that reduce CO_2_ (or formate); 2) acetoclastic methanogens that ferment acetate to CH_4_ and CO_2_; and 3) methylotrophic methanogens that oxidize and reduce methanol to CH_4_ and CO_2_ [14]. All methanogens found in human intestines so far are hydrogenotrophic [15]:

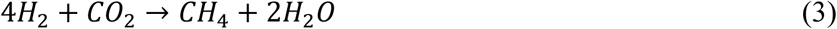

Unlike homoacetogens and SRB, hydrogenotrophic methanogens do not have alternative metabolic pathways: They rely solely on H_2_ and CO_2_ for their metabolism, and detected CH_4_ is always produced via a H_2_-consuming process.

CH_4_ also is unique among the hydrogenotrophs because it is only made by archaea and is not metabolized by the human body [16]. In contrast, H_2_S is produced from other microbial sulfur metabolisms [17] and metabolized by mitochondria in host colonocytes [18]. Likewise, acetate is produced through fermentation by many bacteria [19], and it is a common substrate for any respiring bacteria [20].

Some evidence supports that methanogens have an impact on host metabolism in mouse models [11]. For example, mice inoculated with methanogens showed increased weight and adiposity despite consuming the same amount of food as controls [21,22]. However, the role of methanogens in human metabolism is complicated and controversial. Studies have found conflicting correlations between methanogens and obesity/leanness [23] and anorexia [24].

In this exploratory analysis, we investigated relationships between methanogens and the other hydrogenotrophs, along with the impact of methanogens on human-host metabolism, in samples from a tightly controlled randomized crossover feeding study with well characterized human participants [25]. Briefly, Corbin et al. [25] evaluated the microbial contribution to human-host energy balance using two distinctly different diets: the Western Diet (WD) and the Microbiome- enhancer Diet (MBD). The WD was comprised of foods that were low in fiber and resistant starch, small in particle size (such as peanut butter vs. whole nuts on the MBD), and included processed foods. The MBD, in contrast, was designed to deliver more microbial substrates to the colon by being less absorbable by the host. The MBD contained more whole foods, fiber, resistant starch, and was limited in processed foods. They found that diet altered microbiome’s structure, and that the microbiome’s metabolic activity contributed to host metabolizable energy (ME).

The results from Corbin et al. [25] are particularly well-suited for a deeper investigation into methanogens and their role in the microbiome and host metabolism. On the one hand, that study strictly accounted for energy input (diet), energy expenditure, and energy output (urine, feces, gas). On the other hand, they continuously measured CH_4_ production during the inpatient portion of the study using a first-in-human method within a whole room calorimeter [26].

Combining the study’s CH_4_ measurements with its multi-omic data allows us to detect relationship among the methanogens, the other hydrogenotrophs, and the human host.

Since the hydrogenotrophs compete for H_2_, we hypothesized that high methanogenic activity would lead to a lowering of homoacetogenic and sulfate-reduction activities. These changes in the microbial community would then be associated with factors related to the human’s metabolism: e.g., fecal SCFA output, serum SCFA concentration, and host ME.

## Methods

### Overview of Clinical Study

Details of the clinical study (NCT02939703) from which the data and samples for this work were derived were previously published [25,26]. Briefly, the clinical study was approved by the AdventHealth Institutional Review Board and conducted at AdventHealth Translational Research Institute in Orlando, Florida. After a health screening, 17 participants (9 men and 8 women) completed the study, which was a randomized crossover-controlled feeding study with the WD as a control and the MBD as an intervention. This design minimized the impact of confounders as each participant served as their own control. The study period took place over 61 days. Each participant’s caloric requirement was determined during the baseline period (days 1-9), and meals were prepared uniquely for each participant to maintain energy balance. Participants consumed those meals outpatient for 11 days then inpatient for the next 11 days with a >14-day washout between diet periods. During the 11-day inpatient stay for each diet, each participant’s energy expenditure was measured in whole room calorimeter for 6 days and fecal samples were collected.

### CH_4_ Measurements

CH_4_ release was measured continuously during each 6-day period that the participant was in the whole-room calorimeter. CH_4_ concentration was measured with an off-axis integrated-cavity output spectroscopy (OA-ICOS). A detailed description and validation of the method can be found in Carnero et al. [27].

### Colonic Transit Time

Colonic transit time (CTT) was measured while participants were in the whole-room calorimeter. Participants ingested a SmartPill™ (Medtronic) that sent data to a sensor worn by the participants recording temperature, pressure, and pH [26].

### Host Metabolizable Energy (ME)

Briefly, host ME, the energy from the diet available to the host [28], was computed as the total energy intake minus the energy lost in the feces. Calculation details are published Corbin et al. [25].

### Fecal and Serum Short Chain Fatty Acids

Fecal and fasting serum SCFAs were quantified by targeted metabolomics (Metabolon, Inc., Mooresville, NC). Fecal SCFAs were normalized as published in Corbin et al. [25].

### Quantification of Key Hydrogenotroph Genes

The abundances of the homoacetogens, SRB, and methanogens were measured using the quantitative polymerase chain reaction (qPCR) for genes that identify each hydrogenotroph: *acsB* for homoacetogens, *dsrA* for SRB, and *mcrA* for methanogens. Primers, thermocycler settings, standards, and references for each gene are summarized in **Supplementary Table 1**. All qPCR assays were performed on a Thermofisher Applied Biosystems Quant Studio 3.

Standards for each gene were custom IDT gBlocks gene fragments of the target genes from a representative microbe of each hydrogenotroph group. 7-point calibration curves for each assay were generated in triplicate using gene copy numbers ranging from 10^1^ to 10^8^ of their respective gene standards.

The qPCR values were transformed from log to exponential values, normalized to daily fecal output, giving us daily fecal copy number (gene copy numbers/day). Since each hydrogenotroph contains one gene copy per cell of their respective gene [29–31], qPCR measurements of gene copy number provided an accurate estimate of hydrogenotroph cell numbers/day in the feces.

### DNA Sequencing and Sequence Processing

DNA sequencing and subsequent sequence processing were performed as previously described in Corbin et al. [25].

### RNA Sequencing

Fecal-sample processing, RNA extraction, library prep, and mRNA sequencing were performed at the University of North Carolina at Chapel Hill Microbiome Core (Chapel Hill, NC, USA), which is supported by the following grants: Gastrointestinal Biology and Disease (CGIBD P30 DK034987) and the UNC Nutrition Obesity Research Center (NORC P30 DK056350). RNA was extracted using the Qiagen RNeasy PowerMicrobiome Kit (Cat No./ID: 26000-50). RNA depletion was performed using QIAseq FastSelect –5S/16S/23S Kit (Cat No./ID: 335925), and library prep was performed using QIAseq Stranded Total RNA Lib Kit (Cat No./ID: 180745).

Samples were sequenced using the Illumina HiSeq 4000 PE150 platform. To avoid batching effects, fecal samples were randomized prior to nucleic acid extraction and all samples were sequenced at the same time.

### RNA-Sequence Processing

RNA-sequencing outputs were quality controlled with FastQC (version 0.12.0) [35]. Adapters were trimmed using TrimGalore (version 0.6.5) [36], RNA sequences were aligned against Hg38 (GRCh38.p14) using STAR (version 2.7.11a) [37], aligned sequences were removed, and reads were then paired and annotated using HUMAnN3 (version 3.8) [38] using standard parameters. A small pseudo-count (equal to half of the lowest non-zero count) was added to any zeros and transcript abundances were centered log-ratio transformed as suggested for the analysis of compositional data [39,40].

### Wilcoxon Signed Rank Test

The Wilcoxon signed rank test was used to compare the median log_10_(daily fecal copy number/day) for hydrogenotroph marker genes *mcrA* (methanogens), *acsB* (homoacetogens), and *dsrA* (SRB). Samples in which a gene was undetected were given a gene copy number of 1 and then log transformed.

### Categorizing High and Low Methane Producers

Study participants were categorized as either high or low CH_4_ producers. First, methane production rates were log_10_ transformed and then tested for multimodality by the excess mass test (Excess mass = 0.22, p-value < 2.2e-16) using the modetest function from the R package “multimode” (version 1.5) [41]. Next, the threshold for high CH_4_ production was set at 37 mL CH_4_/day using the locmodes function. Participants were evaluated on a per sample basis and categorized as a high or low methane producer irrespective of diet.

### Linear Mixed Model

We used a linear mixed model (LMM) to test the relationship between high and low CH_4_ producers, our primary independent variable of interest, and outcome variables. We accounted for the effects of colonic transit time, diet, diet period, and diet sequence by including them as covariates in our LMMs. We included participant ID as a random factor to account for the non-independence of the samples. Details about dependent and independent variables, covariates, and random term used in our LMMs can be found in **Supplement Table 1**. LMMs were run using the lmer command from the R package “lmerTEST” (version 3.1-3) [42].

The residuals of each LMM were evaluated for a normal distribution by using qqnorm and shapiro.test commands, both of which were from the R package “stats” (version 4.2.2) [43]. If model residuals were non-normal a transformation was applied. Fecal and serum SCFAs were square-root transformed. Because host ME was negatively skewed (or left-tailed), the ME values were reflected before they were transformed, meaning each ME value was subtracted from a constant (the maximum ME value plus 0.1) and then log transformed [44]. The Benjamini-Hochberg method was used to correct for multiple comparisons and adjusted p-values ≤0.10 were considered significant. The same analyses were completed with CH_4_ as a continuous variable. The results of the continuous-variable analyses are in the Supplementary Data.

### Network Analysis

Microbiome network analysis was performed using the Cytoscape (version 3.10.2) [45] plugin Conet (version 1.1.1) [46] an ensemble co-occurrence analysis tool. Conet was run with default settings using Pearson, Spearman, Mutual Information, Bray-Curtis dissimilarity, and Kullback-Leibler dissimilarity to detect associations between microbes. The Benjamini-Hochberg method was used to correct for multiple comparisons and adjusted p-values ≤0.5 were considered significant.

## Results and Discussion

### Homoacetogens and SRB were present in all participants, but not all participants had methanogens

We first investigated the abundance of methanogens, homoacetogens, and SRB among the trial participants. We performed qPCR targeting genes that encode enzymes in the hydrogenotrophic pathways for each group: *mcrA* for methanogens*, acsB* for homoacetogens, and *dsrA* for SRB.

We detected methanogens in only 9 participants for both diets and one participant for only the MBD (**Figure 2**). Consequently, the average log_10_(gene copy number/day) was 5.3±4.8 for methanogens. In contrast, we detected homoacetogens and SRB in all 17 participants for both diets. On average, homoacetogens were more abundant than SRB: log_10_(gene copy number/day) of 10.2±0.46 for homoacetogens versus 8.8±0.86 for SRB, but homoacetogens and SRB were more abundant than methanogens. Detecting methanogens in only some participants is consistent with past results showing that the abundance of *M. smithii* in the human gut was either high or very low [47,48]. The presence or absence of methanogens can have important implications for hydrogenotrophs and the microbiome. In mixed-culture experiments, SRB became the dominant hydrogenotroph when methanogens were inhibited [11]. In co-culture experiments, *Bacteroides thetaiotaomicron* expressed more glycoside hydrolases, enzymes that break down polysaccharides, and degraded and fermented more glycans when the methanogen *M. smithii* was present, compared to when the SRB *D. piger* was present [49].

**Figure 2.**
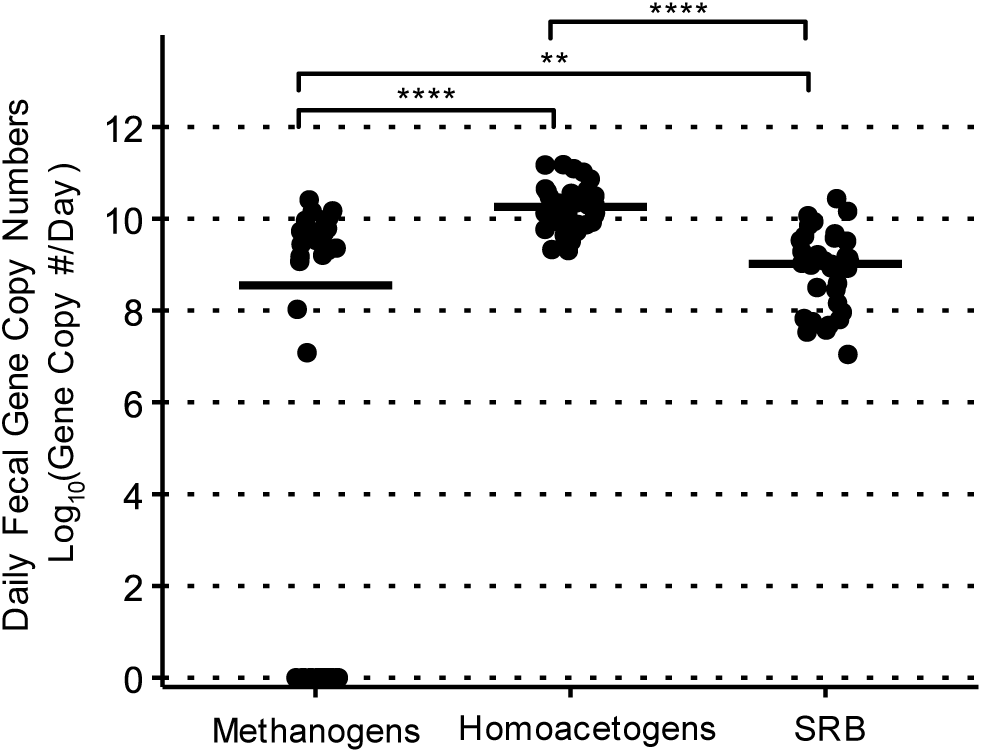
Abundances of hydrogenotrophs in daily fecal copy numbers (log_10_ (gene copy/day)) measured by qPCR. Homoacetogens and SRB were detected in all participants in both diets. Methanogens were only detected in 9 participants in both diets and one participant in the MBD. Homoacetogens were more abundant than SRB and methanogens. SRB were more abundant than methanogens. Abundance medians are marked by the horizontal black bars. Comparison tests between abundance medians were performed with Wilcoxon signed rank test. *: adj. p-value <0.10, **: adj. p-value<0.05, ***: adj. p-value<0.01, ****: adj. p-value<0.0001. NS is non-significant with an adjusted p value > 0.10.

### Homoacetogens and SRB did not show patterns of competition with methanogens

The absence of methanogens in a portion of the participants might lead to differences in competition for H_2_ among the hydrogenotrophs. We hypothesized that, in the absence of methanogens, homoacetogens or SRB (or both) would become more abundant and active. To test this hypothesis, we compared mean methanogen, SRB, and homoacetogen daily fecal copy numbers and gene and transcript abundances between high and low CH_4_ producers High CH_4_ producers had higher methanogen daily copy numbers in the feces and the abundance of *mcrA* genes and transcripts, compared to low CH_4_ producers (**Figure 3A**). In contrast, homoacetogen daily copy numbers, *acsB* genes, and *acsB* transcripts were not significantly different between groups (**Figure 3B**). While SRB copy numbers were higher in high CH_4_ producers, *dsrA* gene and transcript abundances were not significantly different between groups (**Figure 3C**). The trends were similar when this analysis was done using CH_4_ production as a continuous variable (**Figure S1**).

**Figure 3.**
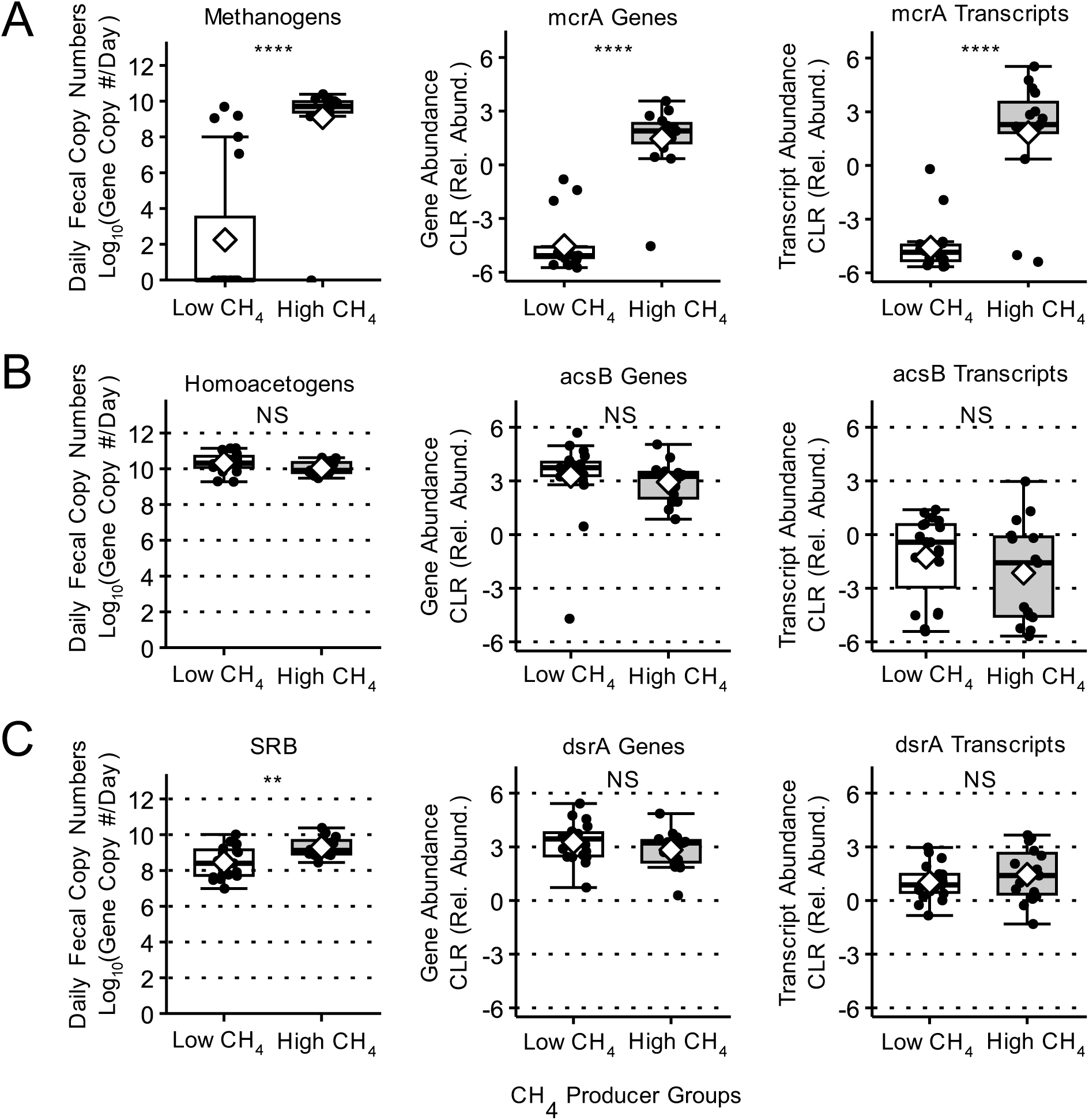
Comparisons of mean hydrogenotroph daily fecal copy numbers, gene and transcript abundance between CH_4_ groups. A) Methanogen daily fecal copy numbers, mcrA genes, and mcrA transcripts were significantly higher in high CH_4_ producers than low CH_4_ producers. B) Homoacetogen daily fecal copy numbers, acsB genes, and acsB transcripts were not significantly different between CH_4_ groups. C) Sulfate reducing bacteria (SRB) daily fecal copy numbers were significantly higher in high CH_4_ producers than low CH_4_ producers, but dsrA genes, and dsrA transcripts were not significantly different between CH_4_ groups. CLR is the centered log-ratio transformation. Group means are marked by a white diamond. Group medians are marked by horizontal black line. All means comparisons were made by linear mixed model. *: adj. p-value <0.10, **: adj. p-value<0.05, ***: adj. p-value<0.01, ****: adj. p- value<0.0001. NS is non-significant with an adjusted p value > 0.10.

These results suggest that competition for H_2_ was not a factor limiting the growth and accumulation of SRB and homoacetogens. One explanation is that SRB and homoacetogens have diverse metabolisms and do not rely solely on H_2_ as an electron donor [50,51]. For example, SRB can utilize lactate for sulfate respiration [17] or for fermentation in the absence of sulfate [52,53]. Homoacetogens can also ferment sugars instead of oxidizing H_2_ during hydrogenotrophic homoacetogenesis [54]. Although homoacetogens can grow mixotrophically, using hydrogenotrophy and fermentation simultaneously for energy generation, an experiment evaluating the mixotrophy of *Blautia coccoides*, a homoacetogen found in the human gut, revealed that hydrogenotrophy was inhibited by glucose [55]. The high abundance of *acsB* genes relative to the low abundance of *acsB* transcripts suggests that homoacetogens may not have been relying on H_2_ for electrons. A second explanation is that the electron flow to CH_4_ was too small to have had a major impact on the other hydrogenotrophs’ metabolic function: Electron flow from COD ingested to CH_4_ was approximately 1% of the electron-equivalent intake to the large intestine (Davis et al. 2024).

### High CH_4_ producers had higher serum propionate concentrations than low CH_4_ producers

Although the hydrogenotrophs did not appear to compete for H_2_ and the flow of electrons into CH_4_ was small, CH_4_ production still may have had an impact on the H_2_ concentration in the large intestine. Because fermentation is inhibited by high H_2_ partial pressure [56], additional H_2_ consumption by methanogens may still have relieved a thermodynamic inhibition on fermentation and led to greater fecal SCFA outputs and serum SCFA concentrations. Fecal SCFA were not significantly different between groups, although the means were consistently lower for the high CH_4_ producers (**Figure 4A**). In contrast, serum propionate concentrations were higher in high CH_4_ producers than low producers (**Figure 4B**). Total fecal SCFAs, fecal acetate, fecal propionate, and serum propionate were significantly positively associated with CH_4_ when these analyses were done using CH4 as a continuous variable (**Figures S2**). Isotope studies have shown that serum propionate is mostly microbially produced in the intestine [20,57]. Therefore, the opposing relationships of fecal SCFA and serum propionate with CH_4_ production suggest a faster rate of SCFA uptake in participants that produced more methane.

**Figure 4.**
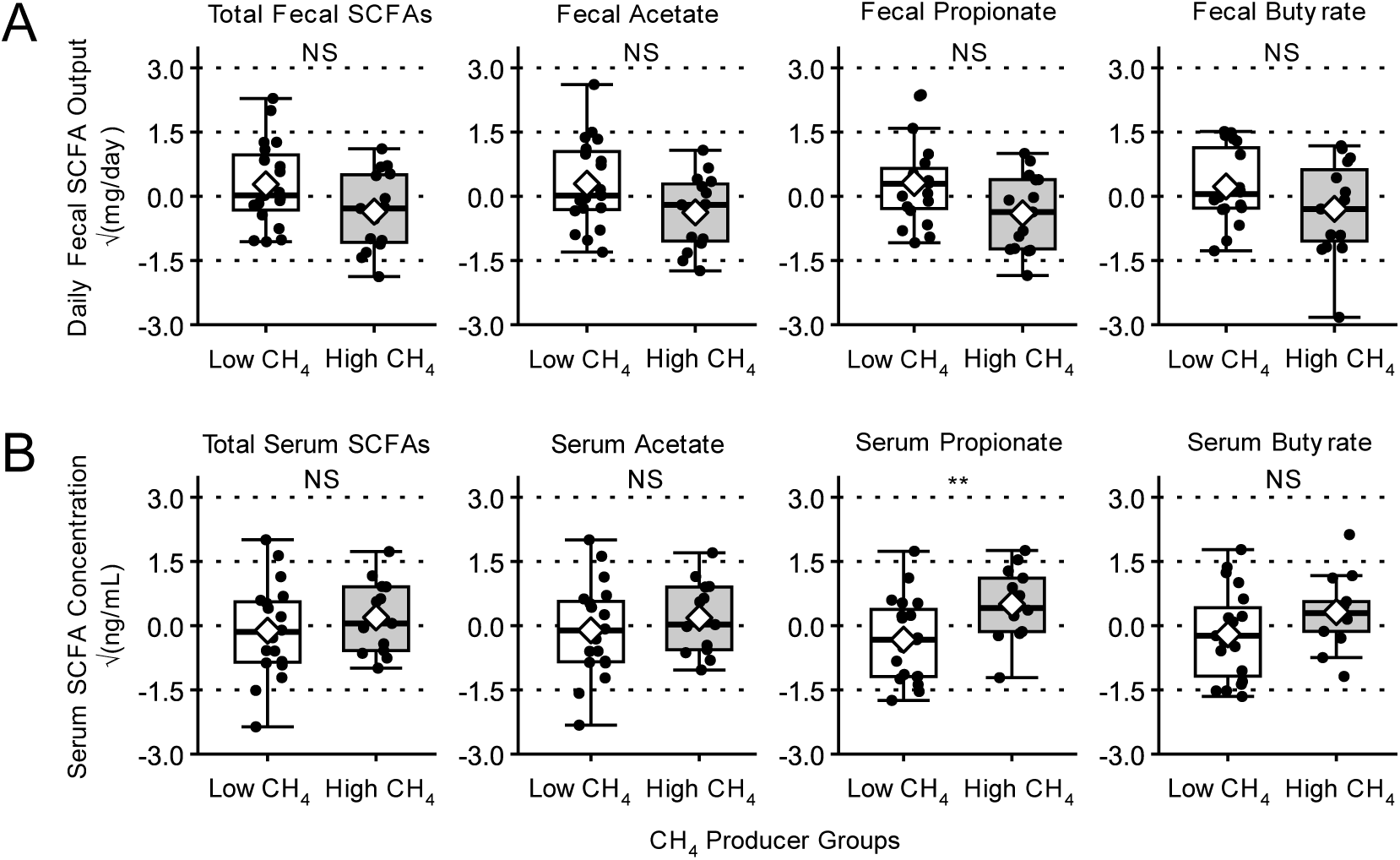
Comparisons of mean fecal SCFA output and serum SCFA concentration between CH_4_ groups. A) Total fecal SCFA, fecal acetate, fecal propionate and fecal butyrate were not significantly different between CH_4_ groups. B) Serum propionate concentrations were significantly higher in high CH_4_ producers than low CH_4_ producers, but total serum SCFA, serum acetate, and serum butyrate were not significantly different between CH_4_ groups. Fecal SCFA output and serum SCFA concentrations were square root transformed. Group means are marked by a white diamond. Group medians are marked by horizontal black line. All means comparisons were made by linear mixed model. *: adj. p-value <0.10, **: adj. p-value<0.05, ***: adj. p-value<0.01, ****: adj. p-value<0.0001. NS is non-significant with an adjusted p value > 0.10.

### High CH_4_ producers had a higher abundance of gene and transcripts for a propionate-producing pathway than low CH_4_ producers

Given the positive association between serum propionate and CH_4_-production rate, we investigated if propionate production could have been altered by methanogenesis. We evaluated the relationship between the CH_4_-production rate and the abundance of key microbial genes and transcripts for microbial pathways for propionate production. The common propionate-producing pathways, the succinate and acrylate pathways [58], consume H_2_ [59]. The succinate pathway has two variants, one using methylmalonyl-CoA decarboxylase (*Mmd*) and the other using methylmalonyl-CoA carboxyltransferase (*MMCT*). The acrylate pathway is characterized by the acryloy-CoA reductase (*Acr*) [58,60]. The abundances of the *Mmd* genes and transcripts were higher in high CH_4_ producers (**Figure 5A**), *MMCT* genes and transcripts were not different between groups (**Figure 5B**), and *Acr* gene abundance, but not transcript abundance, was higher in high CH_4_ producers (**Figure 5C**). Similar results were obtained when this analysis was done using methane as a continuous variable (**Figure S3**).

**Figure 5.**
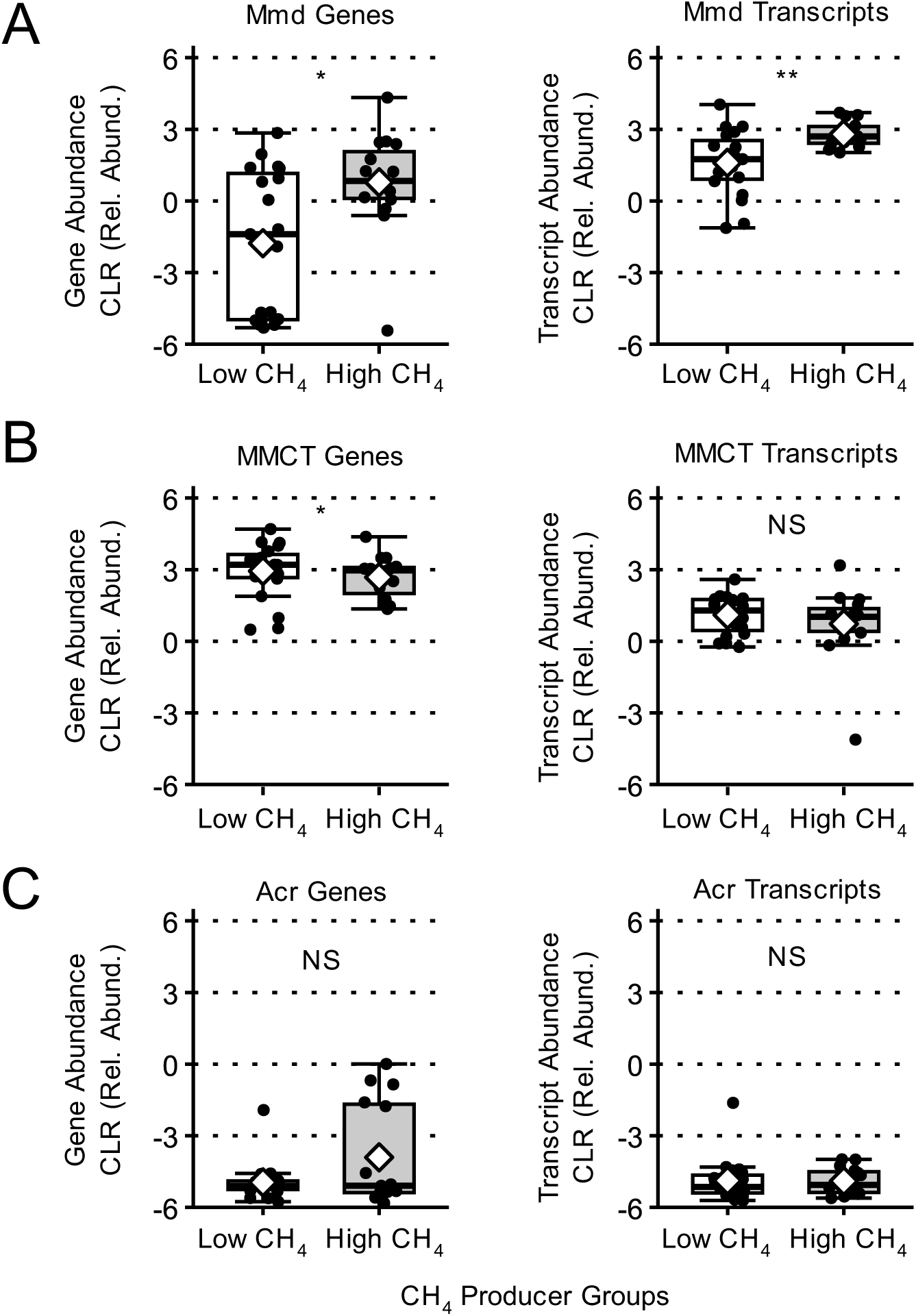
Comparison of gene and transcript abundance for the propionate-producing succinate (Mmd and MMCT) and acrylate (Acr) pathways between CH_4_ groups. A) Mmd (methylmalonoyl-CoA decarboxylase) gene and transcript abundance were significantly higher in high CH_4_ producers than low CH_4_ producers. B) MMCT (methylmalonyl-CoA carboxyltransferase) gene and transcript abundance were not significantly different between high and low CH_4_ producers. C) Acr (acroyl-CoA reductase) genes were significantly higher in high CH_4_ producers than low CH_4_ producers, but transcripts were not significantly different between high and low CH_4_ producers. CLR is centered log-ratio transformation. Group means are marked by a white diamond. Group medians are marked by horizontal black line. All means comparisons were made by linear mixed model. *: adj. p-value <0.10, **: adj. p-value<0.05, ***: adj. p-value<0.01, ****: adj. p-value<0.0001. NS is non-significant with an adjusted p value > 0.10.

Miceli et al. [61] showed that methanogenic microbial communities, while maintaining CH_4_ production rates, increased propionate production in response to increased carbohydrate availability for fermentation. We calculated the average amount of H_2_ consumed by methanogenesis and propionate production in participants with detectable methanogen fecal copy numbers. The calculations can be found in the Supplementary Materials. We found that methanogenesis consumed on average around 0.12 e^-^ eq /day of H_2_, whereas we estimated that propionate production could have consumed as much as 716 e^-^ eq/day of H_2_, assuming all propionate was produced via the succinate pathway. Our results and calculations support the notion that electron flow to propionate production was substantially greater than to methanogenesis.

### Host ME was greater in high CH_4_ producers

Absorption of microbially generated SCFA can contribute up to 10% of daily caloric uptake [25,62], and our results indicate that high CH_4_ production may be an indicator or biomarker of increased SCFA absorption by the host. Thus, we compared host ME between high and low CH_4_ producers. High CH_4_ producers had a significantly higher host ME than low producers **(Figure 6A**). When the analysis was performed with CH_4_ as a continuous variable (**Figure 6B**), the relationship was significantly positive for all participants on both diets, and each diet showed a similar correlation, although all ME values were higher for the WD than the MBD. Because methanogens in the human colon comprise only around 1.2% of the microbiome [63] and rely solely on H_2_ plus CO_2_ or formate for energy [64,65], they neither produce nor consume SCFAs. In principle, methanogens could have increased host ME by consuming H_2_ and thermodynamically accelerating fermentation to SCFAs. However, as we showed above, the electron flow to CH_4_ was small relative to the potential electron flow to propionate. A more compelling explanation is that methanogens were a key component of a microbial community that enabled greater host uptake of energy from the large intestine.

**Figure 6.**
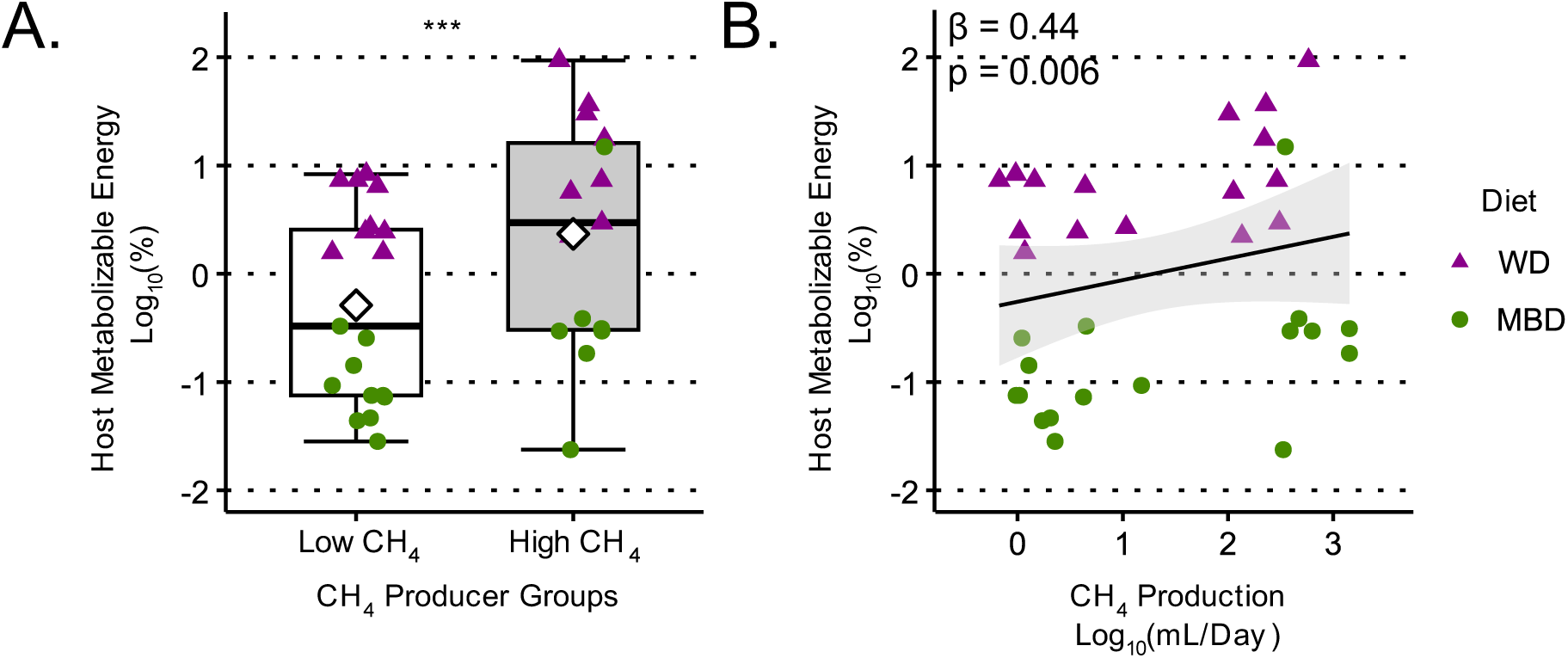
Host ME in high and low CH_4_ producers and by CH_4_ production rate. A)Host ME was significantly higher in high CH_4_ producers than low CH_4_ producers. B) Host ME was significantly positively associated with CH4 production. β is the standardized effect size for the linear mixed model describing the relationship between Host ME and CH_4_ production. WD is the western diet. MBD is the microbiome enhancer diet. Group means are marked by a white diamond. Group medians are marked by horizontal black line. All means comparisons and variable relationships were made by linear mixed model. *: adj. p-value <0.10, **: adj. p-value<0.05, ***: adj. p-value<0.01, ****: adj. p-value<0.0001. NS is non-significant with an adjusted p value > 0.10.

### Fiber-degrading bacteria and propionate-producing bacteria co-occur with *M. smithii*

Considering the positive association between host ME and CH_4_-production rate, along with methanogens’ inability to degrade fiber or ferment sugars, we looked for co-occurring bacteria that made the methanogenic microbial community better equipped to degrade and ferment organic substrates. We used CoNet [46], a network co-occurrence inference tool, to find bacteria strongly associated with *Methanobrevibacter smithii*, the predominant methanogen detected in our samples. After correction for multiple comparisons, 19 bacteria were positively correlated, and 8 bacteria were negatively correlated with *M. smithii* (**Figure 7**).

**Figure 7.**
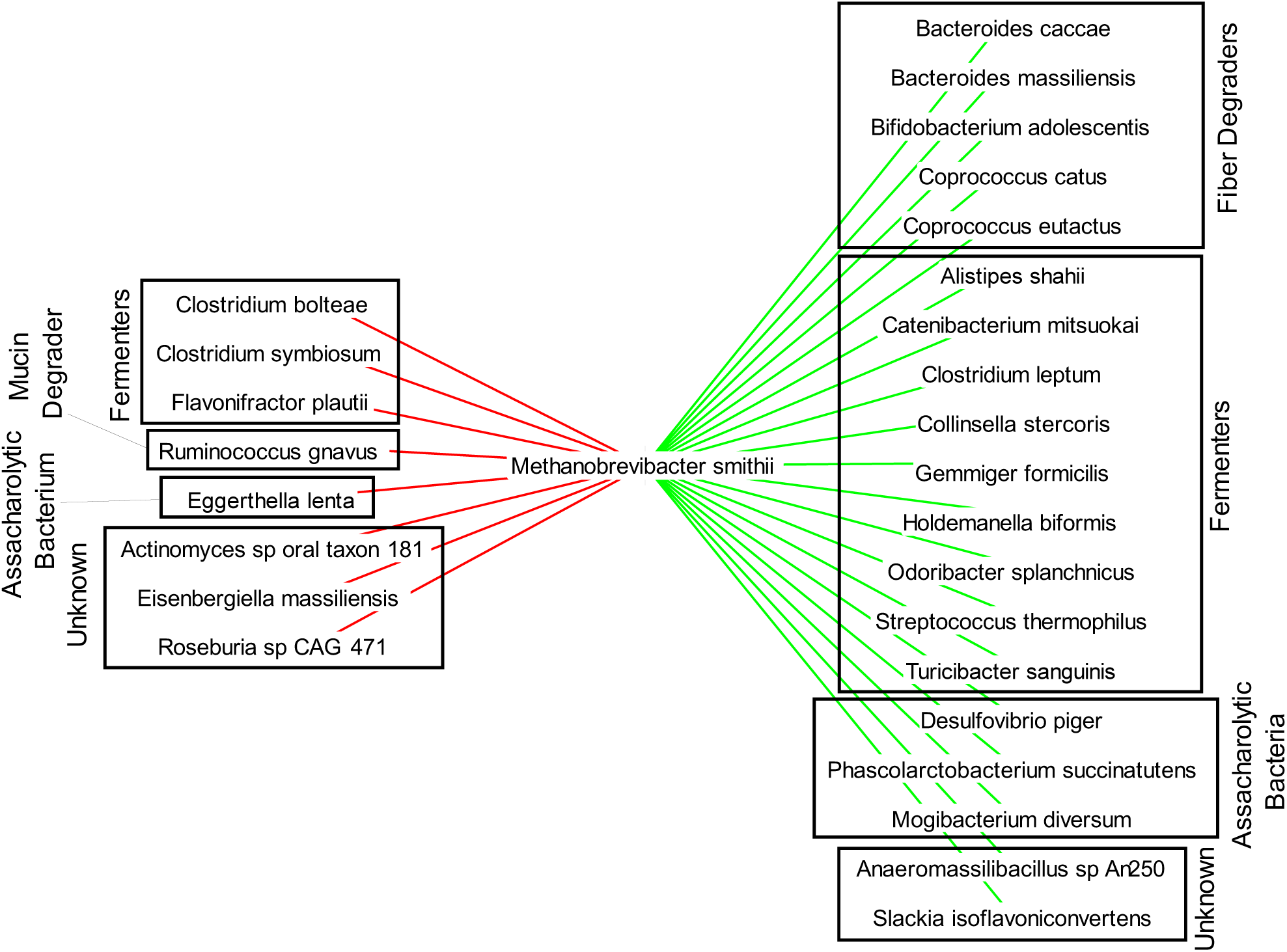
Network from co-occurrence analysis of M. smithii and associated bacteria. Red lines represent negatively associated bacteria, and green lines represent positively associated bacteria. Asaccharolytic bacteria lack the ability to degrade sugars and utilize other substrates such as amino acids or cross-feed on bacterial metabolites such as lactate. All relationships were statistically significant after correcting for multiple comparisons (adj.p < 0.05).

Bacteria positively associated with *M. smithii* were either fiber degraders -- *Bacteroides caccae* [66]*, B. massiliensis* [67]*, Bifidobacterium adolescentis* [68]*, Coprococcus catus* [69], and *C. eutactus* [70]) -- or a diverse group of fermenters -- (*Alistipes shahii* [71]*, Catenibacterium mitsuokai* [72]*, C. leptum* [73]*, Colinsella. stercoris* [74]*, Holdemanella biformis* [75],

*Odoribacter splanchnius* [76], and *Streptococcus thermophilus* [77]). Four of the 19 positively associated bacteria produce propionate via the succinate pathway (*B. caccae, B. massiliensis, O. splanchnicus,* and *Phascolarctobacterium succinatutens* [78]), which aligns with the results on propionate-producing pathways presented in **Figure 5**.

Bacteria negatively associated with *M. smithii* were fermenters -- such as *Clostridium bolteae* [79]*, C. symbiosum,* and *Flavonifactor plautii* [80] *--* mucin-degraders -- such as *Ruminococcus gnavus* [81] and asaccharolytic bacteria -- or bacteria that cannot metabolize sugars, such as *Eggerthela lenta* [82]. All these bacteria, except *E. lenta,* can produce acetate and either ethanol or lactate. This ability is important because it allows these bacteria to continue fermentation despite high H_2_ partial pressure by shifting production from acetate to less reduced ethanol or lactate [6,83]

We interpret that a network of fiber degraders, fermenters, propionate producers, and methanogens provided the host with more SCFA and contributed to higher host ME (**Figure 6**). **Figure 8** shows how the bacteria positively associated with *M. smithii* fit in a hypothesized trophic chain. Fiber degraders broke down polysaccharides into sugars; fermenters consumed the sugars to release acetate, butyrate, propionate, and H_2_; and methanogens and propionate producers consumed the H_2_ and kept fermentation thermodynamically favorable. Thus, the methanogenic microbial communities could utilize more complex substrates than non-methanogenic microbial communities; this explains how the methanogenic communities could produce more SCFAs for the host to absorb.

**Figure 8.**
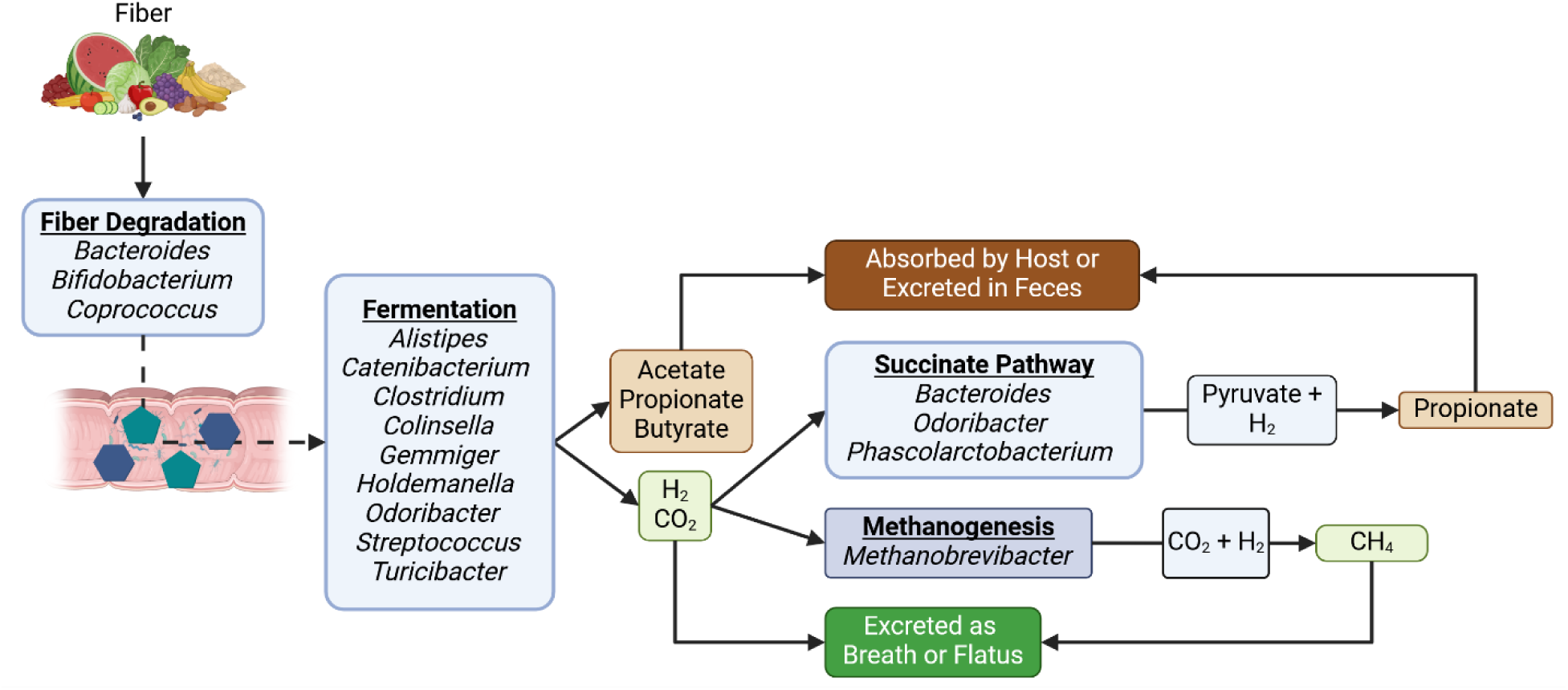
Trophic chain formed in a methanogenic gut microbiome. The bacteria positively associated with M. smithii form a trophic chain that efficiently degrades and ferments fiber and effectively removes excess H_2_ produced through fermentation by methanogenesis and the propionate-producing succinate pathway. Figure created using Biorender.

## Conclusion and Future Directions

Our results show that methanogens and methane production were present in only about one-half of the study participants. While methanogen fecal copy numbers and *mcrA* genes and transcripts were higher in high CH_4_ producers, SRB and homoacetogen fecal copy numbers, genes, and transcripts had no relationship with CH_4_ production. Consequently, we did not see evidence that methanogens’ uptake of H_2_ to produce methane affected the other hydrogenotrophs. This probably occurred because SRB and homoacetogens, unlike methanogens, do not rely on H_2_ oxidation for energy generation. Methanogenesis was associated with higher host ME and serum propionate, but not with fecal SCFAs, suggesting that methanogens were linked to enhanced SCFA uptake. We also found that bacteria co-occurring with methanogens are well-suited to degrade and ferment fiber, as well as consuming H_2_ through propionate production. Taken altogether, our results add important missing mechanistic insights into the relationship between methanogens and obesity. Methanogens appear to be part of a microbial consortium capable of enhanced energy extraction and absorption.

Future research should focus on investigating whether methane or methanogens affect production and absorption of SCFAs in the colon. As we show, the impact that methanogens have on host metabolism may not be in removing H_2_, but rather by influencing other physiological parameters, such as SCFA production and possibly SCFA absorption. Alternatively, methanogens might simply constitute a detectable signal for enhanced energy extraction.

Understanding the interactions of methanogens with the human host and other microorganisms in the human large intestine could lead to dietary or other means to modulate methanogen activity in ways that improve the host’s metabolic health. Evaluating the presence and activity of methanogens that accumulate on the intestinal epithelium would provide important information that cannot be obtained solely from fecal samples. Additionally, future experiments that involve *in vitro* microbial communities of homoacetogens, SRB, and methanogens would allow for direct measurements of all hydrogenotrophic activities for a wide range of conditions; this would provide a higher resolution picture of the structural and functional relationships among the hydrogenotrophs.

