## Supplementary Data for "Methanogens are associated with altered microbial production of short-chain fatty acids and human-host metabolizable energy"

**Supplementary Material**

**Supplementary Table 1: qPCR primer and themocycler details**

| **Group** | **Gene** | **Primers** | **Thermocycler Settings** | **Reference** | **Standards from** |
| --- | --- | --- | --- | --- | --- |
| Homoacetogens | *acsB* | ACS_f  ACS_r | 95°C 10 s, 52°C 20 s, 72°C 30 s, 40 cycles | [32] | *Blautia hydrogenotrophica*  DSM 10507 |
| Sulfate-reducing bacteria | *dsrA* | DSR1-F DSR-R | 95°C 10 s, 60°C 60 s, 35 cycles | [33] | *Desulfovibrio piger*  isolate FI11049 |
| Methanogens | *mcrA* | mlas  mcrA-rev | 95°C 30 s, 55°C 45 s, 72°C 30 s, 40 cycles | [34] | *Methanobrevibacter smithii*  ATCC 35061 |

| **Supplementary Table 2. Linear Mixed Model Variables** |
| --- |
| Independent Variables (IV): |
| ·         CH4 Producer Group (high and low) |
| Outcome Variables (OV): |
| ·         Total fecal SCFAs normalized to PEG recovery (μg/day) |
| ·         Fecal acetate normalized to PEG recovery (μg/day) |
| ·         Fecal propionate normalized to PEG recovery (μg/day) |
| ·         Fecal butyrate normalized to PEG recovery (μg/day) |
| ·         Total Serum SCFAs (ng/mL) |
| ·         Serum Acetate (ng/mL) |
| ·         Serum Propionate (ng/mL) |
| ·         Serum Butyrate (ng/mL) |
| ·         Host Metabolizable Energy (%) |
| ·         Gene copy number of key hydrogenotroph genes |
| ·         Gene transcript abundance of key hydrogenotroph genes |
| Covariates: |
| ·         Randomized diet assignment |
| ·         Diet Period |
| ·         Diet Sequence |
| ·         Colonic Transit Time (CTT) |
| Random Factor: |
| ·         Participant ID |
| ·         Accounts for non-independence of the samples |

**Calculations for daily average hydrogen consumption for methanogenesis and the succinate pathway**

We calculated the average amount of H2 consumed by methanogenesis and propionate production in participants with detectable methanogen fecal copy numbers. We assumed that all propionate production went through the hydrogen-consuming succinate pathway and that 95% of microbially produced propionate is absorbed in the colon and 5% of microbially produced propionate remains in the feces [83].

| **category** | **constant** | **value** | **unit** |
| --- | --- | --- | --- |
| standard conditions | pressure | 1 | atm |
|  | universal gas constant | 0.08206 | (L atm)/(K mol) |
|  | temperature | 298 | K |
| Stoichiometry (equations below) | H_2_/CH_4_ stoichiometry coefficient | 4 | mol H_2_/ mol CH_4_ |
|  | H_2_/C_3_H_6_O_2_ stoichiometry coefficient | 2 | mol H_2_/ mol C_3_H_6_O_2_ |
| molecular weight | propionate molecular weight | 74.079 | g C_3_H_6_O_2_ / mol C_3_H_6_O_2_ |
| assumption 2 | fraction of produced propionate in feces | 0.05 | g C_3_H_6_O_2_ feces / g C_3_H_6_O_2_ total |

methanogenesis stoichiometry

$$4H_{2}+CO_{2}\to CH_{4}+2H_{2}O$$

succinate pathway propionate production stoichiometry

$$C_{3}H_{4}O_{3}+ 2H_{2} \to C_{3}H_{6}O_{2}+ H_{2}O$$

| **category** | **calculated variable** | **value** | **unit** |
| --- | --- | --- | --- |
| methane | daily avg methane production volume | 365.13 | mL CH_4_ / d |
|  | daily avg methane production mols | 0.01 | mol CH_4_/ d |
|  | daily avg hydrogen consumption mols | 0.06 | mol H_2_ / d |
| fecal propionate | daily avg fecal propionate mass | 121.33 | mg C_3_H_6_O_2_ feces / d |
|  | daily avg propionate production mass | 2.43 | g C_3_H_6_O_2_ / d |
|  | daily avg propionate production mols | 179.76 | mol CH_6_O_2_ / d |
|  | daily avg hydrogen consumption mols | 359.51 | mol H_2_ / d |

Linear mixed model analyses with CH4 production as a continuous variable.

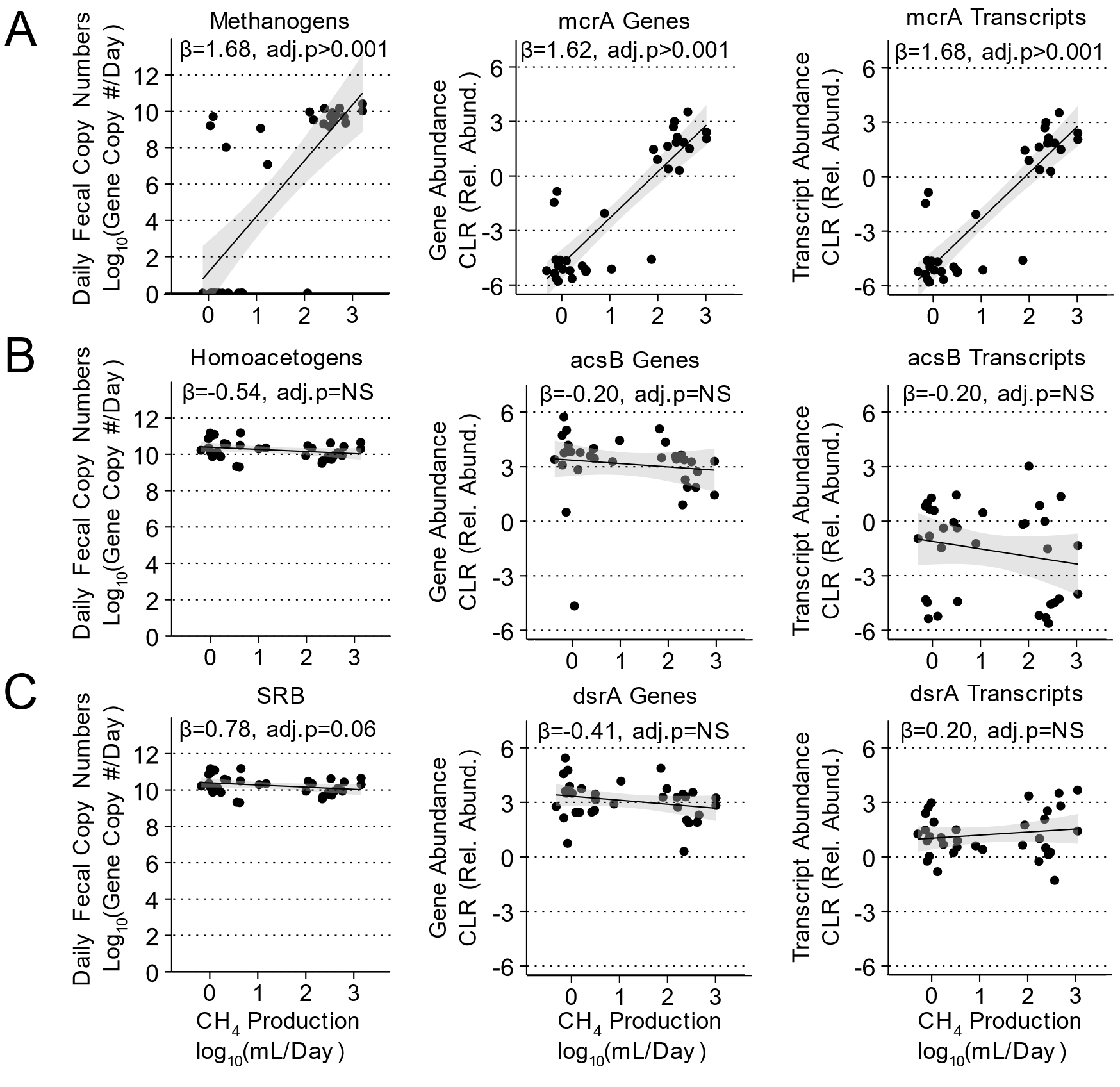

**Figure S1. Hydrogenotroph daily fecal copy numbers, gene and transcript abundance, and their relationship with CH_4_-production rates.** A) Relationship of methanogen daily fecal copy numbers, mcrA genes, and mcrA transcripts with CH_4_ production. B) Relationship of homoacetogen daily fecal copy numbers, acsB genes, and acsB transcripts with CH_4_ production. C) Relationship of sulfate reducing bacteria (SRB) daily fecal copy numbers, dsrA genes, and dsrA transcripts with CH_4_ production. β is the standardized effect size for the linear mixed model describing the relationship between CH_4_ production and daily fecal copy numbers, gene relative abundances, or transcript relative abundances. NS is non-significant with an unadjusted p value > 0.05.

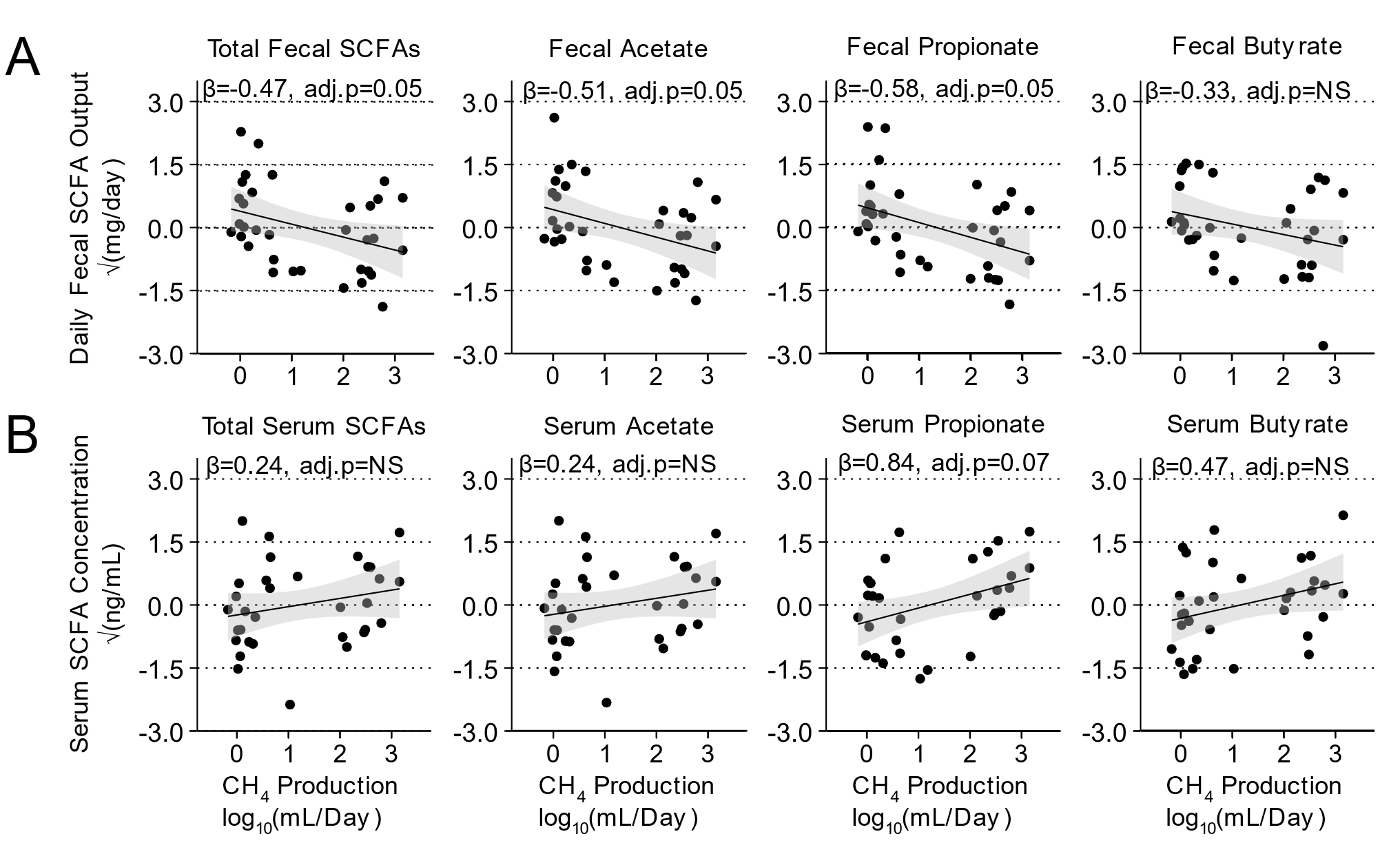

**Figure S2. Relationships of fecal SCFA output (mg/day) and serum SCFA concentration (ng/mL) to the CH_4_-production rate (mL/day).** A) Total fecal SCFA, fecal acetate, and fecal propionate were negatively correlated with the CH_4_-production rate. B) Serum propionate concentrations were positively associated with the CH_4_-production rates. β is the standardized effect size for the linear mixed model describing the relationship between CH_4_ production and fecal SCFA output and serum SCFA. NS is non-significant with an unadjusted p value > 0.05.

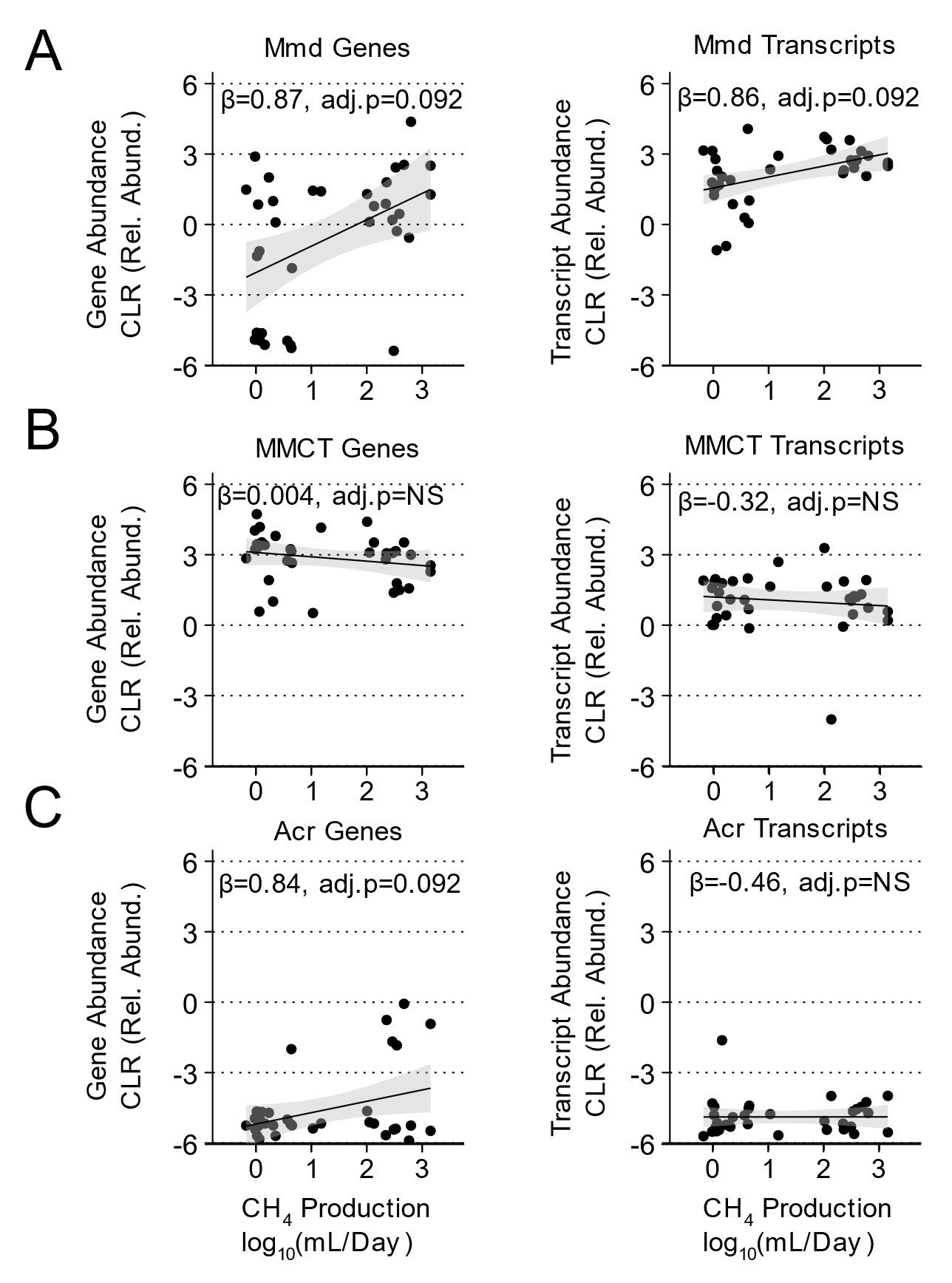

**Figure S3. Gene and transcript abundance for the propionate-producing succinate (Mmd and MMCT) and acrylate (Acr) pathways by CH_4_ production.** A) Association of Mmd (methylmalonoyl-CoA decarboxylase) gene and transcript abundances with CH_4_ production. B) Association of MMCT (methylmalonyl-CoA carboxyltransferase) gene and transcript abundance with CH_4_ production. C) Association of Acr (acroyl-CoA reductase) gene and transcript abundances with CH_4_ production. β is the standardized effect size for the linear mixed model describing the relationship between CH_4_ production and key gene and transcript abundances of the propionate-producing succinate and acrylate pathways. NS is non-significant with an unadjusted p value > 0.05.
